# Acoustic community structure and seasonal turnover in tropical South Asian birds

**DOI:** 10.1101/518985

**Authors:** Anand Krishnan

## Abstract

Birds produce diverse acoustic signals, with coexisting species occupying distinct ‘acoustic niches’ to minimize masking, resulting in overdispersion within acoustic space. In tropical regions of the world, an influx of migrants from temperate regions occurs during winter. The effects of these migrants on acoustic community structure and dynamics remain unstudied. Here, I show that in a tropical urban bird community, the influx of winter migrants is accompanied by a turnover of the acoustic community. However, in spite of this turnover, the acoustic community remains overdispersed in acoustic niche space. The winter acoustic community additionally exhibits lower frequency-band diversity, consistent with species singing less continuously, as well as lower phylogenetic diversity. My data thus suggests that acoustic niches and community structure are stable across seasons in spite of species turnover. Migrants occupy similar regions of acoustic space as residents, and are relatively closely related to some of these species. Their arrival therefore leads to greater phylogenetic clustering in the winter, and thus lower phylogenetic diversity, although the acoustic community remains overdispersed. Studying seasonal dynamics of acoustic communities thus provides valuable insight into assembly processes, as well as a potential framework for long-term monitoring of urban ecosystems.

## Introduction

The diverse communication functions of the songs of breeding birds include territorial advertisement and attracting mates (1). In the more well-studied temperate regions of the world, many birds sing during the spring and summer breeding season, migrating southward during the winter months (2). Within communities of vocal species (or acoustic communities), simultaneous vocalizations may mask each other, posing a barrier to efficient communication (3–6). As a result, acoustic signals of a community of birds (as well as other diverse animals) may exhibit overdispersion (7–9), occupying distinct regions of acoustic parameter space (or “acoustic niches”, for example singing at different times or frequencies) to minimize overlap (10–17). The development and expansion of passive acoustic recording and monitoring techniques has enabled study of the acoustic space of entire bird communities (10,18–20), however, much remains to be understood about the dynamics of these communities. In particular, the influence of seasonal dynamics (21,22) on singing activity and acoustic community structure has received relatively little study.

Studies of the seasonal dynamics of bird song are particularly relevant in tropical landscapes, which are home to the largest proportion of the world’s biodiversity. Owing to the high species diversity of the tropics, and impending threats from deforestation and climate change, tropical acoustic communities provide valuable insights informing both our understanding of behavioral ecology and conservation (23). In Neotropical forest bird communities, signals of coexisting birds may diverge as a result of competition, and the overall temporal patterns of singing activity are driven by community composition (7,8). In addition to the highly diverse resident breeding avifauna, tropical regions also receive a large influx of winter migrants from temperate regions, which may increase the local species diversity (24,25). Many of these migrants remain highly vocal on their wintering grounds, and their effect on the structure and dynamics of tropical acoustic communities has not, to my knowledge, been studied. The influx of winter migrants into acoustic niche space, many of which are close relatives of tropical resident species, could result in more dense ‘packing’, and thus clustering in acoustic space (26,27). Alternatively, resident species may drop out of the acoustic community (this means an absence of vocalizations, either implying local movements or becoming silent), thus resulting in species turnover of the acoustic community. In this latter scenario, overdispersion of the acoustic community, and thus acoustic niche structure, may be predicted to persist within the winter. However, these patterns and seasonal dynamics remain poorly known.

Here, I quantify seasonal community dynamics of avian vocal activity in an urban dry deciduous scrub-grassland in peninsular India. This habitat is highly biodiverse, generally underrepresented in studies of tropical ecology (28), and under severe threat due to extensive destruction and land-use change (29). Using recordings of the acoustic community across wet (monsoon) and dry (winter) seasons, I quantify whether the acoustic community exhibits overdispersion consistent with the presence of acoustic niches, and whether the influx of winter migrants alters this community structure (i.e. the distribution of species within acoustic space) or instead results in turnover of the acoustic community. Finally, I examine whether the arrival of migrants alters the phylogenetic (and thus species) diversity of the acoustic communities. Because scrub habitats possess a varied and conspicuous avifauna, most of which breed in the monsoon (30), and also receives a large influx of winter migrants (31), studying this habitat provides valuable insight into seasonal patterns in biodiversity.

## Materials and Methods

### Study Site

I conducted this study in the Vetal Tekdi Biodiversity Park, a small (<80ha) remnant patch of tropical dry-deciduous scrub-grassland mosaic within the city of Pune, Maharashtra, India. The typical vegetation of this area includes native vegetation such as *Acacia* and *Anogeissus*, and a largely open savanna-type habitat structure with a grassy understory (29) (Figure 1). The study site is a public park at the top of a hill at about 790m asl. The slopes of this hill have been planted over by non-native trees, and native vegetation now only remains in higher areas. This habitat exhibits pronounced seasonality, with the Southwest monsoons between June and September, and also undergoes annual burning of grass (32,33). Using multiple microphones, it was possible to collect simultaneous and comprehensive information on bird vocalizations across the landscape. Additionally, it was possible to identify vocalizing species and assign calls to them relatively easily in this open habitat, which is key to identifying species vocalizations in large passively recorded datasets. The avifauna of this park includes several species characteristic of scrub landscapes across India, for example *Francolinus sp*, *Lanius sp* and *Pericrocotus erythropygius* (31,34) (see Supplementary Data), and may thus be considered representative of these habitats.

**Figure 1:**
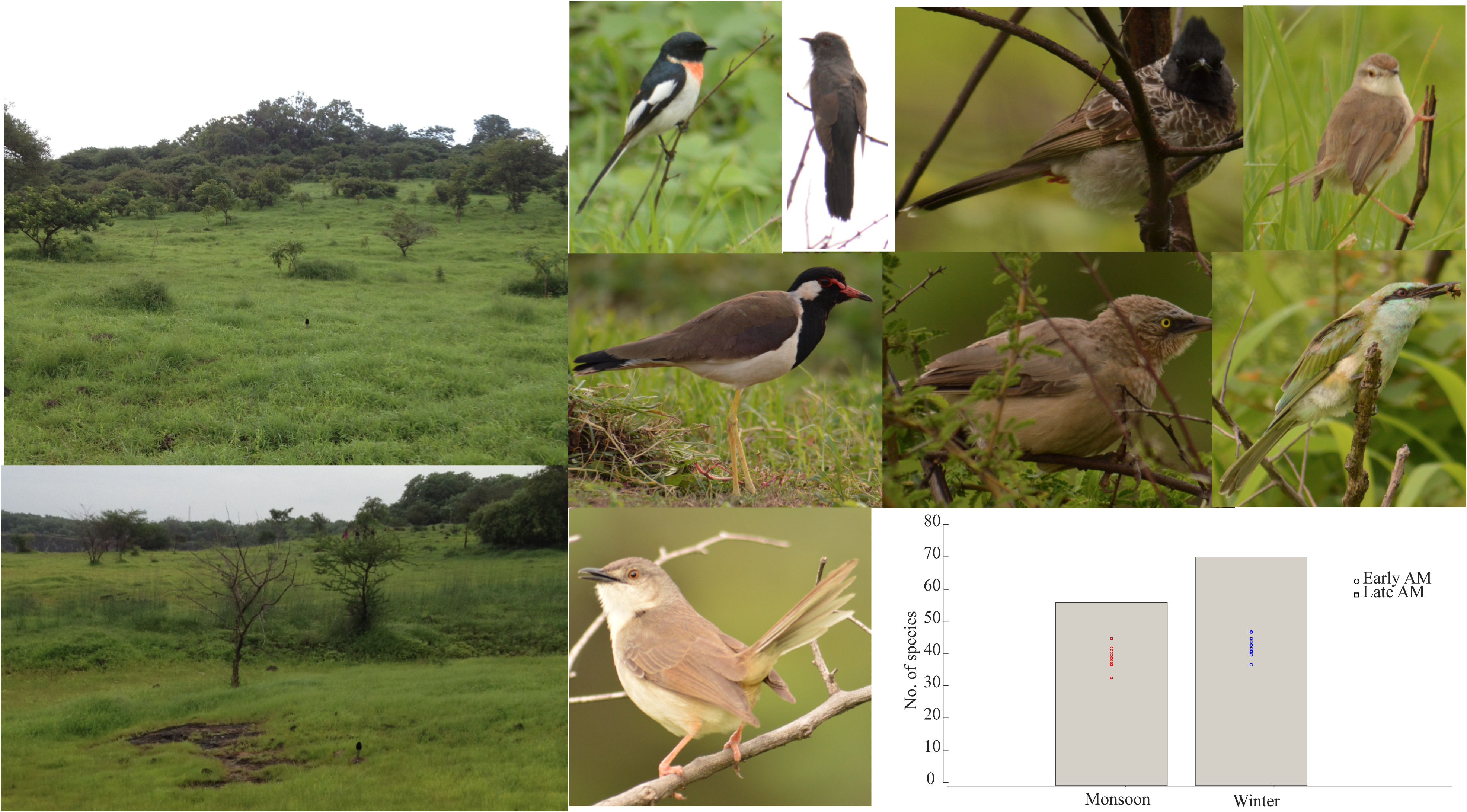
The study site, Vetal Tekdi Biodiversity Park, and some birds found in the study area. All images by Vivek Kannadi, Science Media Center, IISER Pune. The graph represents total number of species recorded in each season, and each point represents the number of species recorded in each individual 45- minute recording.

### Recordings

In order to quantify seasonal changes in bird singing activity during the dawn chorus, I conducted recordings during August 2017 (the monsoon breeding season for many resident birds), and November-December 2017 (winter migrant season). The recording apparatus I used consisted of four Sennheiser (Wedemark, Germany) ME62 omnidirectional microphones connected to a Zoom H6 (Tokyo, Japan) portable recorder for simultaneous, synchronized recordings. The four microphones were connected to the recorder using 10m long XLR cables, each leading off to the north, south, east and west of the recorder. The microphones were then placed on the ground to record singing activity. Using four sensitive, low-noise omnidirectional microphones enabled me to maximize detection of high-frequency species with soft calls, and thus simultaneous, comprehensive coverage of all bird species vocalizing within this landscape. Additionally, this allowed me to minimize differences in detection due to microphone placement, or due to differences in temperature and humidity across seasons, because four microphones increase the likelihood of detecting species whose signals might be affected by these factors. Because I did not observe spatial heterogeneity in singing activity within this rather open habitat, I performed pooled analyses of all four microphone channels except where otherwise indicated. On each sampling day, I recorded 45 minutes of bird singing activity between 630 and 730AM (the hour of sunrise, referred to as the early morning recording), and another 45 minutes between 730 and 830AM (referred to as the late morning recording) (broadly following a similar sampling strategy to (8)). In total, the primary dataset consisted of six sampling days in the monsoon and six in the winter. At 90 minutes a day, this resulted in a total of 18 sampling hours of bird song activity, or 72 hours of audio data in total across four microphones. Before and after the data collection periods, I made recordings and surveys of bird vocalizations at the site using a single microphone, both to aid identification of calls detected during the census and as a reference library to calculate call parameters.

### Species activity and seasonal turnover

To determine relative activity patterns of each species in the acoustic community across seasons, I divided each 45-minute recording into five-minute segments. Within each five-minute segment, I identified all species of vocalizing bird across all four microphone channels (pooled) both by ear and by spectrographic visualization of calls in Raven Pro 1.5 (Cornell Laboratory of Ornithology, Ithaca, NY, USA). Most bird species in this habitat are familiar and common Indian birds, including in urban environments, and are thus easily identifiable by voice, as well as by comparison to reference recordings I made earlier. Therefore, although there is a remote possibility of mis-detection in any census using a human observer, this is very unlikely to alter patterns of presence-absence, particularly of common species. I constructed a presence-absence matrix where a 1 indicated presence of a species’ vocalizations and 0 their absence in each 5-minute sample. Once this had been carried out for the entire dataset (12 45-minute recordings per season, and four microphone channels per recording), I proceeded to examine seasonal patterns for species that were present in 10% or more of total 5-minute samples in each season (8) (which I henceforth refer to as the acoustic community), to avoid bias introduced by vocally “rare” species. For these species, I determined what I henceforth refer to as an acoustic “abundance index”. In order to account for non-independence of consecutive 5- minute samples, I drew one 5-minute sample at random for each species from each 45-minute recording (thus a total of 12 per species per season, six from early morning recordings and six from late morning recordings), and calculated the percentage of 1’s (or presences) in this random draw (schematic of workflow in Figure 2A). The process was repeated 10000 times for each species for each season. The average percentage of presences across 10000 random draws is the acoustic abundance index, which represents the probability of detecting a species’ vocalizations in a randomly-drawn 5-minute acoustic sample. A species that is vocally ubiquitous would score close to a 1 in abundance index, and a rarer species would score a low value. This avoids potential bias introduced by a species being highly vocal on one day and silent on another, by ensuring that samples from each recording day are given equal weight. To calculate species turnover and diversity accounting for abundance, I calculated Jaccard similarity indices between the two communities, and Shannon-Weiner alpha-diversity indices for both monsoon and winter acoustic communities using the vegan (35) package in R (36).

**Figure 2:**
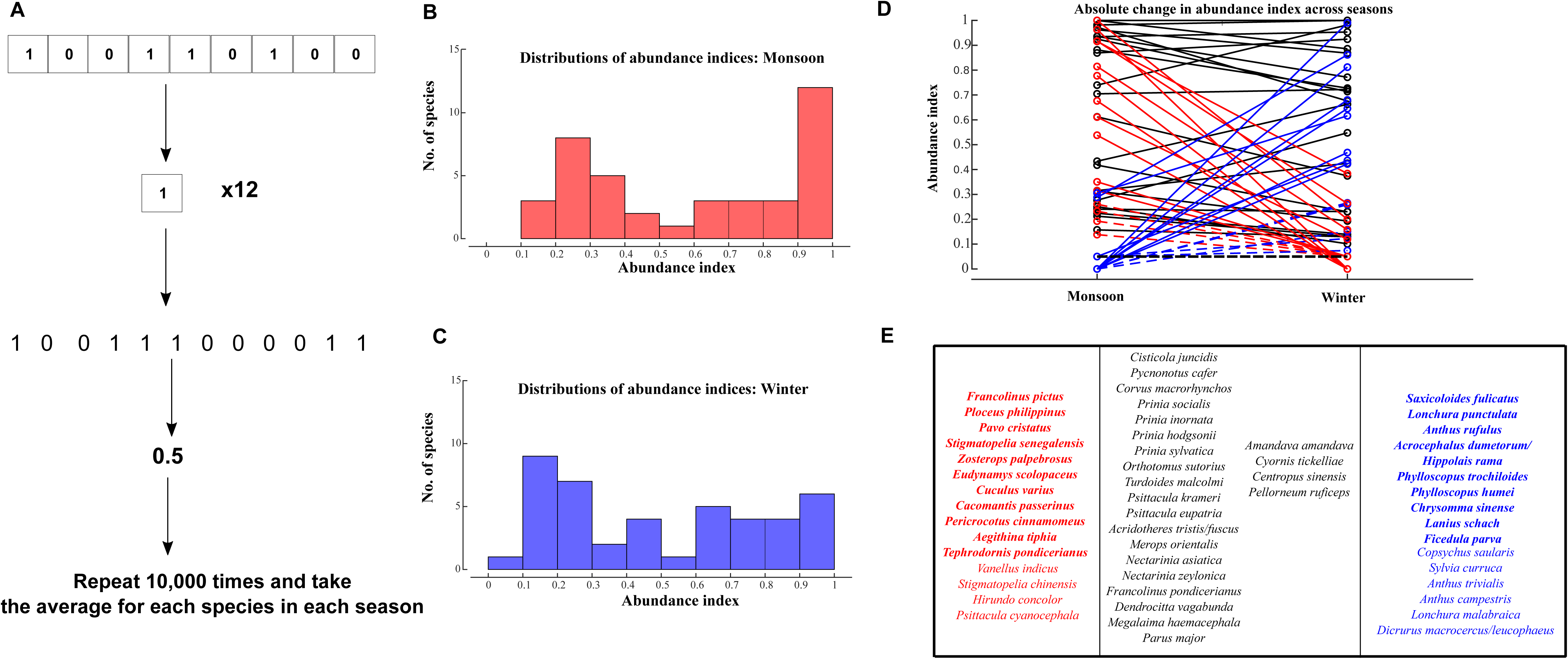
(A) Calculating abundance indices for each species per season. Each 45-minute recording was broken down into 9 5-minute samples, and presence-absence of each species’ vocalizations was indicated by a 1 or 0. One of these 9 samples was drawn at random from each recording, resulting in 12 values per season (six early morning and six late morning recordings). The percentage of 1’s in the resulting 12 values is the acoustic abundance index (in this case 0.5). I repeated this random sampling procedure 10,000 times for each species in each season to obtain the final average value of the abundance index. **(B,C)** Distributions of abundance indices for 40 monsoon (A) and 43 winter (B) species comprising the respective seasonal acoustic communities. **(D,E)** Turnover in species composition of the acoustic community, represented by the seasonal change in abundance index. Species decreasing abundance index in winter by >1 standard deviation are in red solid lines (D) and bold red text (E), and species increasing by the same in winter are the blue solid lines (D) and bold blue text (E). Species that do not exhibit this change are in black in both (D) and (E). Species represented by dashed lines in (D) and regular blue or red text in (E) are those that were detected often enough to calculate an abundance index (although low) in one season but not the other. I also consider these species as exhibiting seasonal turnover.

### Acoustic space and community structure

To further examine seasonal patterns in the diversity of acoustic communities, I used Raven Pro to identify clean examples of calls for each of the species present in the monsoon and winter acoustic communities. I used recordings made outside of the study period from the same site wherever clean examples were not present within the recordings, and for three species, supplemented this with recordings made from peninsular India (as close to Pune as possible, analyses suggested no differences in the parameters calculated) and archived on the Xeno-canto (https://www.xeno-canto.org) and AVoCet (http://avocet.zoology.msu.edu) bird song databases. Using these vocalizations (at least 10 vocalizations per species where possible), I calculated six temporal and frequency parameters in Raven Pro: average peak frequency, maximum and minimum peak frequency, frequency bandwidth (90%), note duration and relative time of peak frequency. I performed a principal components analysis on the correlation matrix of this data to reduce dimensionality. To compare acoustic niche space and structure across seasons, I performed a MANOVA using the manova1 function in MATLAB (Mathworks, Inc, Natick, MA, USA), and one-sample Kolmogorov-Smirnov tests to test for overdispersion of each seasonal community compared to a uniform distribution (9).

### Acoustic diversity indices

Using the first 20 minutes of each 45-minute recording sample (making sure they were free of rain or loud anthropogenic noise), I calculated three indices of acoustic or frequency band diversity (20) using the soundecology (37) package in R to determine overall levels of singing activity. These were the acoustic diversity index (ADI), acoustic evenness index (AEI, where lower values indicate more spread across the frequency spectrum, and thus more evenness), and the bioacoustic index (BI), a measure of acoustic diversity that is more robust against abiotic noise (20,38,39). To quantify changes in singing activity for 10 common species across seasons, I selected the first five minutes from each of six 45-minute recordings for each season (12 in total, free from anthropogenic or weather-related noise). I determined the total percentage of time spent vocalizing across seasons from these five-minute samples (Supplementary Data).

### Phylogenetic diversity indices

Finally, to estimate the phylogenetic diversity of the acoustic communities across seasons as a comparison to their species diversity and seasonal dynamics, I downloaded a phylogeny pruned to contain all the 53 species (including shared species) across both communities from the avian Tree of Life (40). This meta-tree provides 100 different possible phylogenetic hypotheses of the relationships between selected species, and all indices calculated were estimated for each of these 100 trees, thus giving a distribution of 100 index values. I further pruned these trees down from 53 to 40 and 43 species respectively, to calculate separate indices for monsoon and winter acoustic communities. Using these values and the abundance indices for each species, I calculated three commonly measured indices of phylogenetic diversity using the picante (41) package in R: Faith’s phylogenetic diversity (PD) (42), the Mean pairwise phylogenetic distance between species in a community (MPD), and the mean nearest-neighbor phylogenetic distance between species (MNTD) (43) (100 values for each). For the last two, I calculated both raw values and abundance-weighted diversities for both monsoon and winter acoustic communities. To compare phylogenetic diversity values across seasonal acoustic communities, I used paired statistical tests in MATLAB, as each pair of values were calculated using one of the 100 possible phylogenetic trees. For consistency with the phylogenetic analyses, I follow the taxonomy adopted in this tree throughout. Because the acoustic community here is composed of species across multiple families, the broad tree topology does not change much across the 100 possible phylogenies. The distributions of index values reported here are thus robust for broad comparisons across clades.

## Results

### Species diversity of the avian acoustic community

Across the entire dataset, I identified vocalizations of 85 bird species (list of species in Supplementary Data) (44). Of these, 56 species were recorded during the monsoon sampling and 70 species were recorded during the winter, suggesting higher overall species richness during the winter (Figure 1). During the monsoon, early and late morning censuses both recorded an average of 39 species (rounded off) respectively (differences between early and late morning were not statistically significant in a paired t-test: t=0.2571, dF=5, p=0.8074), whereas winter censuses recorded an average of 41 and 44 species in the early and late morning respectively (again not significantly different from each other: t=1.4, dF=5, p=0.2204), per census. Across seasons, I performed unpaired t-tests on early and late morning censuses respectively to determine if more species were recorded in winter. Early morning censuses detected similar numbers of species across seasons (t=1.2677, dF=10, p=0.2336). However, late morning censuses in winter detected significantly more species than in the monsoon (t=2.6451, dF=10, p=0.0245), providing some support for higher overall species richness in the winter. For abundance analyses of the acoustic communities (Figure 2A), however, I selected only species present in >10% of 5-minute samples in each season (see Methods). This resulted in estimates of abundance indices for 40 species in the monsoon and 43 species in the winter (therefore, a total of 53 species in subsequent phylogenetic analyses). Thus, although overall species diversity was higher in the winter, the numbers of regularly vocalizing species comprising the acoustic community were comparable across seasons.

### Seasonal turnover of the acoustic community

The distributions of abundance indices for 40 bird species comprising the monsoon acoustic community and 43 bird species comprising the winter acoustic community are shown in Figure 2B and 2C. Although the number of species in both communities were comparable, this pattern belies a considerable degree of turnover in species composition across seasons. In all, 30 species were shared between the two seasonal communities, resulting in a Jaccard similarity of 0.56. However, if we only consider species above the mean abundance index for each season (roughly 0.5 in both seasons), Jaccard similarity dropped to 0.41, suggesting that 59% of the most abundant species exhibited seasonal changes in acoustic abundance. This is apparent from Figure 2D and 2E, where a number of frequently recorded species (in red) decreased abundance by more than one standard deviation in the winter, whereas others (in blue) increased by more than one standard deviation. I detected resident species such as *Francolinus pictus, Cuculus varius, Cacomantis passerinus* and *Ploceus philippinus* frequently in the monsoon but not at all in the winter, whereas others such as *Stigmatopelia senegalensis, Pavo cristatus* and *Eudynamys scolopaceus* were detected considerably less frequently in the winter. Four monsoon species with low abundance indices (*Vanellus indicus, Stigmatopelia chinensis, Hirundo concolor* and *Psittacula cyanocephala*) were either absent or detected too infrequently in the winter to determine an abundance index. I also list them here under species that decrease abundance in the winter (red dashed lines in Figure 2D). Species increasing in abundance index in the winter included residents such as *Saxicoloides fulicatus* and *Lonchura punctulata* that were also recorded during the monsoon, other residents such as *Lanius schach* and *Chrysomma sinense* that were not, and long-distance migrants such as *Phylloscopus trochiloides*, *Phylloscopus humei* and *Ficedula parva*. Several other species, including the residents *Copsychus saularis* and *Lonchura malabarica*, and the migrants *Sylvia curruca, Anthus trivialis* and *Anthus campestris*, had relatively low winter abundance indices but were either unrecorded or recorded too infrequently to determine an index in the monsoon. They are therefore listed here as species that increase abundance in the winter (blue dashed lines in Figure 2D). Shannon-Weiner diversity indices that account for the abundance (45) (in this case, the abundance indices) of species were comparable for both acoustic communities, at 3.541 for the monsoon acoustic community and 3.559 for the winter, indicating relative stability in acoustic community diversity (i.e. regularly detected species) despite the turnover in species composition.

### Monsoon and winter acoustic communities are overdispersed in acoustic space

The first two principal components of 6 acoustic parameters explained approximately 74% of variation (eigenvalues of 3.35 and 1.07 respectively, see Supplementary Data for values). PC1 exhibited the strongest positive loadings on frequency parameters and weak negative loadings on temporal parameters, whereas PC2 loaded strongly and positively on relative time of peak, and to a lesser extent on note duration. The results of multivariate statistical analysis on these principal components indicate that the overall acoustic parameter (or niche) space occupied by vocalizing species was not significantly different across seasons (Figure 3). MANOVA tests both on the measured parameters and their first three principal components failed to reject the null hypothesis that vocalizations of both monsoon and winter acoustic communities were drawn from the same distribution (On parameters: Wilk’s lambda=0.9331, dF within groups=81, dF between groups-1, total dF=82, p=0.4933, On PCs: Wilk’s lambda=0.9431, p=0.0959). Further, both communities were statistically indistinguishable from a uniform distribution spanning the same range of values in PC1 space (representing the bulk of variation in frequency), which suggests that simultaneously vocalizing birds were overdispersed across acoustic parameter space (One-sample Kolmogorov-Smirnov tests, monsoon: D=0.1268, p=0.5007; winter: D= 0.1673, p=0.1606). This is consistent with species occupying distinct acoustic niches (8,9,46), and with this niche structure remaining stable across seasons, even with the arrival of winter migrants.

**Figure 3:**
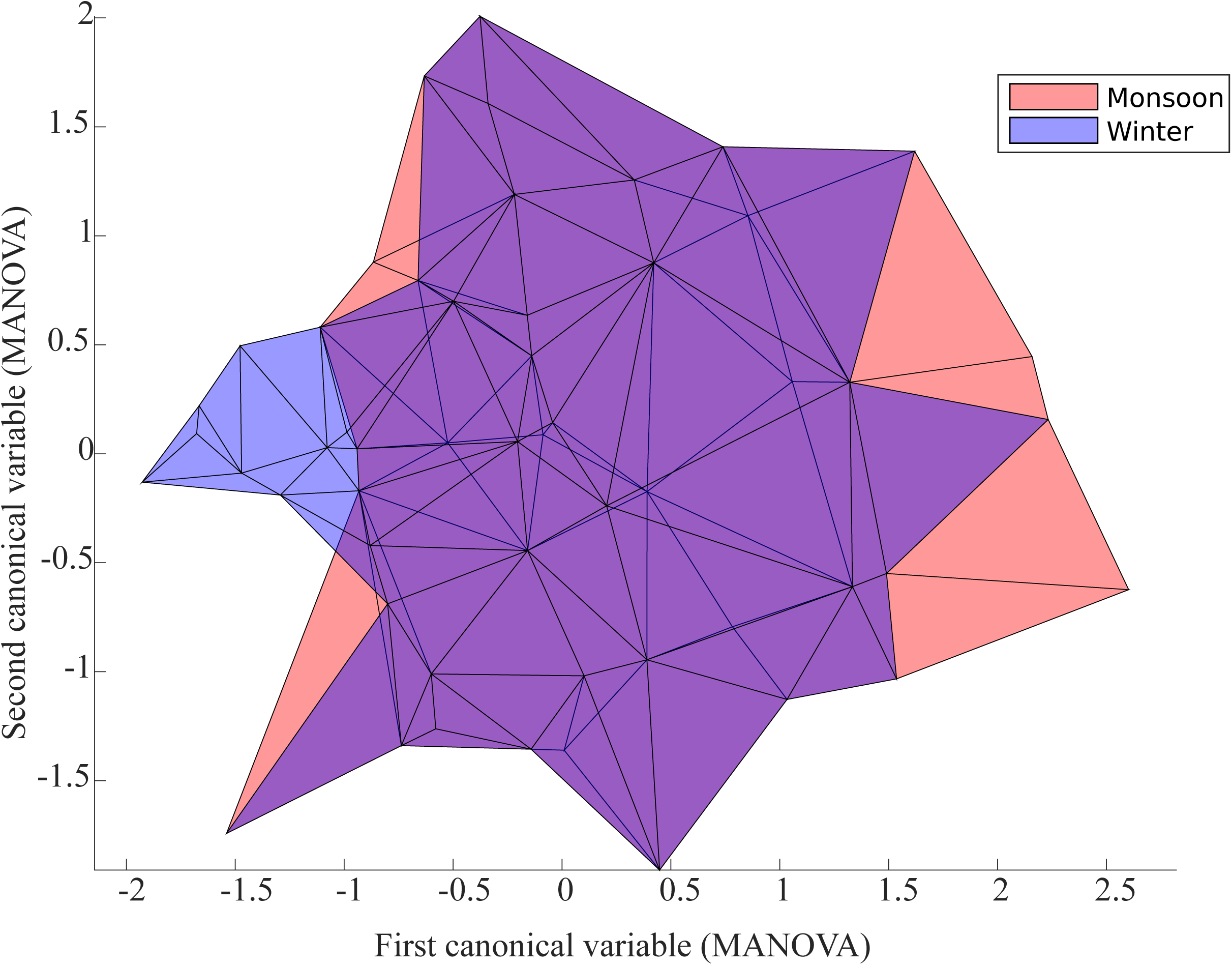
Biplot of first and second canonical variables from MANOVA on the principal components of 6 acoustic parameters, for the monsoon (red) versus winter (blue) acoustic communities.

### Lower frequency band diversity in the winter

Next, I investigated whether turnover and change in species composition was represented by acoustic diversity indices that quantify the relative amount of energy in the frequency band spectrum, and are thus representative of overall levels of singing activity (Figure 4, also see Supplementary Data). All three measures indicate that the monsoon community exhibited greater diversity and evenness than the winter community (Average ADI: Monsoon: 1.4, Winter: 0.59, Average AEI: Monsoon, 0.65, Winter: 0.85, lower values indicate greater evenness, Average BI: Monsoon: 26.65, Winter: 17.11, Wilcoxon signed-rank test, ADI and BI: W=78, N=12/group, P<0.001, AEI: W=0, P<0.001). These values were consistent within each season across recording days, which further indicates robustness across seasons, and that the lower acoustic diversity and evenness observed in winter is genuine. An examination of the relative energy across frequency bands from 0-10 KHz reveals that this reduction appears to be driven by a reduction in activity between roughly 3-7KHz, indicating lower singing activity in the winter(47) (Figure 4B). This suggests two possibilities or a combination of both: firstly, that the species dropping out of the acoustic community in the winter were highly vocal during the monsoon, and their absence drove the lower acoustic indices. Indeed, several species mentioned earlier, with high monsoon abundance indices, were detected less frequently or were absent during the winter. A second contributing factor may be that species that did not change in abundance (and were therefore still frequently vocal) may have been singing shorter bouts outside of their breeding season, reducing the amount of sound in those frequency bands. To test this second hypothesis, I examined changes in percentage of time spent singing for 10 such species over 6 five-minute samples recorded on different days (including all types of vocalizations). For seven of these species, the differences were not statistically significant, but three showed statistically significant (2.59<t<4.01, dF=10, p<0.05, see Supplementary Data) decreases in percentage of time spent vocalizing, even though abundance index does not change (*Prinia socialis, Prinia sylvatica* and *Cisticola juncidis*). These species exhibited high abundance indices across seasons, thus suggesting that they, together with the previously mentioned monsoon vocalizing species, drove the changes in acoustic diversity indices observed here.

**Figure 4:**
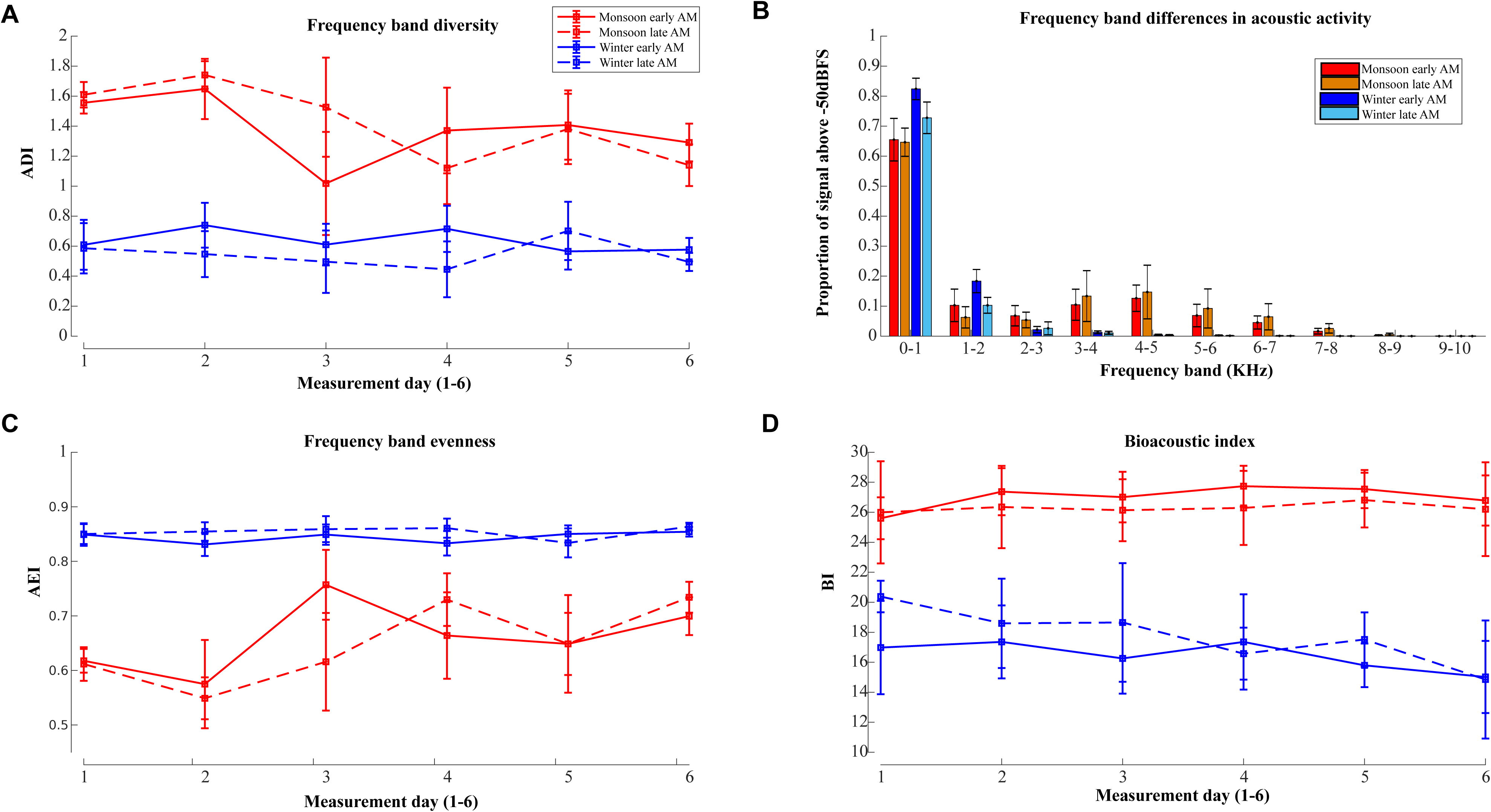
Acoustic diversity indices decline in winter. **(A)** Acoustic diversity index declines in the winter, indicating less diversity of frequency band space. **(B)** The amount of energy across frequency bands suggests that this is driven by a decline in higher frequency activity in winter. **(C)** The Acoustic Evenness Index is higher in winter, indicating lower evenness and more clustering in frequency band space. **(D)** The Bioacoustic Index also supports a decline in higher frequency bands, putatively driven by a decline in continuous singing activity (also see Supplementary Figure).

### Lower phylogenetic diversity in the winter acoustic community

Using a publicly available phylogeny (Figure 5A) (40), I estimated phylogenetic diversity to understand whether the arrival of winter migrants and seasonal turnover influenced the phylogenetic structure of acoustic communities. Three measures of phylogenetic diversity (PD, MPD and MNTD) were all lower for the winter community than for the monsoon (Averages: PD: Monsoon: 1544.83, Winter: 1510,16, MPD: Monsoon: 139.29, Winter: 118.28, MNTD: Monsoon: 50.24, Winter: 48.9). Weighting MPD and MNTD for abundance resulted in slightly lower diversity values across communities but the seasonal trend remained the same (Averages: MPD: Monsoon: 130.92, Winter: 102.13, MNTD: Monsoon: 45.32, Winter: 41.27) (Figure 5B, 5C, also see Supplementary Data). This effect was consistent and statistically significant across all 100 possible phylogenetic trees (PD: paired t-test: t=18.7524, dF=99, p<0.001, effect size=0.38, MPD unweighted: t= 144.18, dF=99, p<0.001, effect size= 3.22, MPD weighted: t=133.44, dF=99, p<0.001, effect size=4.75, MNTD unweighted: t=12.19, dF=99, p<0.001, effect size=0.38, MNTD weighted: t=35.48, dF=99, p<0.001, effect size=1.17), with larger effect sizes after weighting MPD and MNTD for abundance. Thus, although species diversity was comparable in the winter (43 species versus 40 in the monsoon), phylogenetic diversity was lower, indicating more clustering at certain points in the phylogenetic tree of the community.

**Figure 5:**
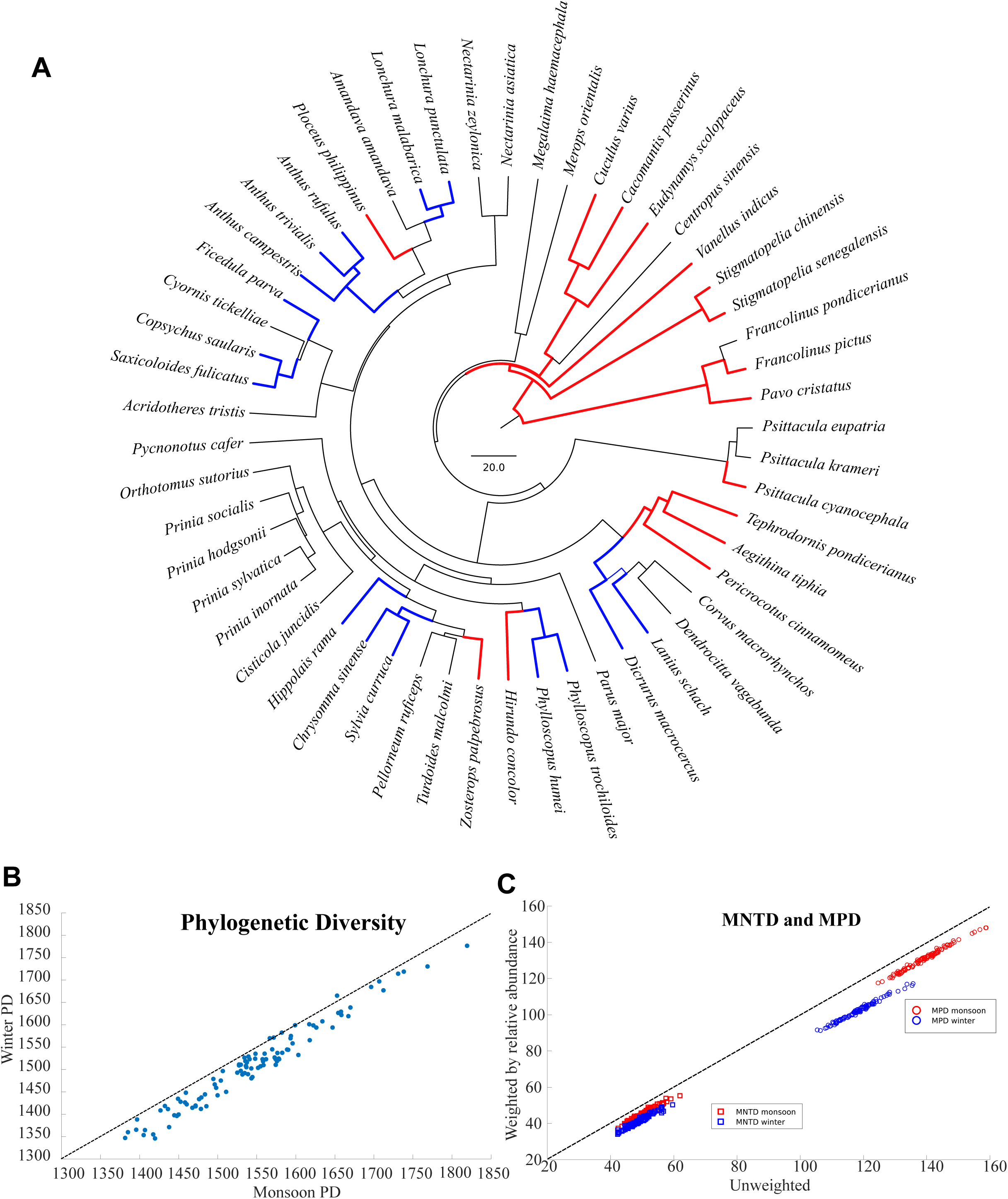
Phylogenetic diversity of the winter acoustic community is lower. **(A)** Maximum clade credibility consensus tree of the 100 possible trees used in the analysis, presented here for illustrative purposes only. Although node support for the consensus tree is low at the tips, the broad species relationships change very little across the 100 trees. Species are colored according to Figure 2 to represent whether they increase or decrease abundance index in the winter. **(B)** Faith’s PD (phylogenetic diversity) is lower in winter than in monsoon, thus lying away from the dotted line, which represents equal values of both. **(C)** Plots of MNTD (squares) and MPD (circles) for monsoon (red) and winter (blue) acoustic communities. The X axis represents unweighted values, and the Y axis the values weighted for abundance. Weighting reduces values of both indices, but the winter community exhibits consistently lower MNTD and MPD than the monsoon community.

## Discussion

To summarize, my study of the avian acoustic community in an urban tropical scrub-grassland habitat uncovers overdispersion of acoustic signals, consistent with the presence of acoustic niches. Although there is considerable seasonal turnover in the species composition of the acoustic community, this overdispersion is maintained across seasons. Additionally, overall use of frequency band space is lower in the winter, putatively driven by a relative lack of continuous vocalizations outside the breeding season (even though detection probabilities of at least some species do not change). Finally, although species diversity in these seasonal acoustic communities is comparable (and late morning winter recordings detected more bird species on average), phylogenetic diversity of the winter acoustic community is actually lower than that of the monsoon. In spite of this increased phylogenetic clustering, species remain overdispersed in acoustic space. Taken together, the data indicate that migrants are close relatives of resident species, and fit into the same acoustic niche space, resulting in a stable, overdispersed acoustic community structure across seasons.

### Bioacoustics and the seasonal dynamics of tropical avian communities

Species in the tropics exhibit considerable seasonal activity changes and local movements, particularly where differences between dry and wet seasons are more pronounced. In semiarid regions and savanna ecotopes of Asia (29), the habitat is relatively open and seasonally dry (32,33). Most resident birds in this habitat breed during the southwest monsoon, and are very vocal between June and October (31). During the winter season, many of the monsoon community’s most vocal resident (‘resident’ is used here to imply birds that do not undertake long-distance migrations) birds fell completely silent, and were also not observed visually during recordings or otherwise (for example, *Cuculus varius*, *Cacomantis passerinus*, *Francolinus pictus* and *Ploceus philippinus*), indicating possible local movements out of the study area. On the other hand, some resident birds (*Lanius schach* and *Chrysomma sinense*, for example) were recorded during the winter only, suggesting local movements into the study area. Other monsoon-breeding birds (*Pavo cristatus* and *Eudynamys scolopaceus*) were recorded during the winter and seen frequently, but were much less vocal. The converse was true of *Saxicoloides fulicatus*, which was detected more frequently in the winter. For these latter species, this may indicate either some local movement, relative silence during one season, or a combination of both. Local movements may be driven by seasonal resources, for example, the seeding of grass (in the case of seed-eaters such as *Lonchura punctulata* and *Lonchura malabarica*) (30,31,34).

In addition, several long-distance migrants were commonly detected in winter and frequently vocal, including *Acrocephalus dumetorum* and *Hippolais rama* (identified visually; it was not possible to distinguish these species acoustically, and they have been treated as a single unit for calculating abundance indices), *Phylloscopus trochiloides, Phylloscopus humei*, and *Ficedula parva*. Thus, a combination of local movements, changes in singing activity and the arrival of winter migrants appears to drive species turnover in the avian acoustic community. This turnover, and the frequent detection of common winter migrants, suggests that acoustic monitoring may provide a valuable tool to detect and monitor the annual influx of winter migrants, as well as to elucidate patterns of local movement in resident species (10,48–50). These local movements remain poorly understood for many bird species, including relatively common ones. Using acoustic methods may provide valuable insight into their movement ecology and behavioral dynamics.

### Stable acoustic community structure across seasons

In spite of the seasonal turnover in species composition, my data suggest that the overall niche structure of bird vocalizations within the community does not change, and the winter community remains overdispersed in acoustic space. This suggests that community structure within acoustic space is stable across seasons, and that long-distance migrants fit into the same acoustic space occupied by resident birds (9,51). This stability across seasons may be a result of migrant species occupying acoustic niches left vacant by the local movements or silence of resident birds. However, the overall use of higher-frequency bands declines in the winter, and indices of frequency-band diversity suggest lower overall singing activity. In addition to the absence of a number of vocal monsoon-breeders, I present preliminary evidence that several of the most vocal resident species (in particular, *Cisticola* and *Prinia* warblers), sing for a greater percentage of time in the monsoon breeding season. A number of birds are known to exhibit higher singing activity coinciding with the breeding season, correlated with hormonal changes such as an increase in male testosterone levels (52). Thus, although the acoustic community remains overdispersed in the winter, niche separation between simultaneously vocalizing species is likely to be of greater importance in the monsoon breeding season, when their vocalizations are more likely to overlap. However, the abundance indices of these species do not exhibit a seasonal change. Therefore, the use of presence-absence abundance indices over 5-minute time blocks may prove useful in quantifying relative abundance regardless of seasonal changes in vocalization. These seasonal changes may, however, result in different acoustic diversity and frequency-band index values even though species diversity is comparable. Acoustic indices (48,49) should therefore be combined with abundance indices to understand community-level seasonal changes in singing activity, at least in tropical bird communities.

### Acoustic niches in tropical bird communities

The acoustic niche hypothesis states that in order to minimize masking or the overlap of sounds, species in a habitat or community may exhibit acoustic signals occupying distinct regions of acoustic space to minimize overlap, and thus masking (10,11,14). In terms of community structure, this is predicted to result in overdispersion of acoustic signals such that they approximate a uniform distribution across acoustic parameter space. Multiple studies have found evidence both for and against this hypothesis in diverse organisms (8,9,11–13,51,53). I find that both monsoon and winter communities are overdispersed in frequency space (PC1 of measured acoustic signal parameters), which accounts for the largest proportion of variation in acoustic signals. This suggests that acoustic niche structure in bird communities is stable across seasons, although, as mentioned above, niche separation may be of greater importance in the monsoon when species vocalize in longer, continuous bouts.

Most migrants recorded during this study, for example, warblers and flycatchers, belong to families that are also represented by resident species (44). All indices of phylogenetic diversity are significantly lower in the winter, even though species diversity is comparable or slightly higher. This indicates that the influx of winter migrants results in more phylogenetic clustering within the acoustic community, and concomitant species turnover while still remaining overdispersed in acoustic space. It is possible that maintaining an overdispersed acoustic niche structure may support the coexistence of multiple closely related species during the non-breeding season. This, however, requires a detailed behavioral study on focal species to address. I note here that my study focuses only on the avian acoustic community within a single urban habitat, to comprehensively understand their seasonal dynamics. However, it is likely that these patterns are general to other habitats receiving an influx of winter migrants as well.

### Conclusions

My study uncovers evidence that bird acoustic communities in tropical deciduous scrubland exhibit distinct niches in acoustic space, such that signals of coexisting species are overdispersed. This acoustic niche structure is stable across seasons to the influx of local and winter migrants, although species composition exhibits turnover within this space, consistent with migrants occupying a similar acoustic space to resident species. Finally, phylogenetic diversity declines in the winter, suggesting that the arrival of migrants results in more phylogenetically clustered communities. A reduction in singing (a proxy of territoriality) by resident birds, and maintaining an overdispersed community structure, may enable their coexistence with multiple closely related migrants, a topic requiring further study. Knowledge of acoustic space and turnover patterns is invaluable to comprehensive assessment, in particular in urban habitats of peninsular India, which are undergoing rapid land-use change. Acoustic monitoring thus serves as a framework for long-term, non-invasive study of the responses of avian communities to land-use and climate change (21,22,49,50).

## Acknowledgments

I thank Rohit Chakravarty and Rashim Malhotra for help with data collection, Raghav Rajan and his lab group for discussions and feedback, Samira Agnihotri and Krishnapriya Tamma for helpful discussions.

